# Complexity Matching and Requisite Variety

**DOI:** 10.1101/414755

**Authors:** Korosh Mahmoodi, Bruce J. West, Paolo Grigolini

## Abstract

Complexity matching emphasizes the condition necessary to efficiently transport information from one complex system to another and the mechanism can be traced back to the 1957 *Introduction to Cybernetics* by Ross Ashby. Unlike this earlier work we argue that complexity can be expressed in terms of crucial events, which are generated by the processes of spontaneous self-organization. Complex processes, ranging from biological to sociological, must satisfy the homeodynamic condition and host crucial events that have recently been shown to drive the information transport between complex systems. We adopt a phenomenological approach, based on the subordination to periodicity that makes it possible to combine homeodynamics and self-organization induced crucial events. The complexity of crucial events is defined by the waiting-time probability density function (PDF) of the intervals between consecutive crucial events, which have an inverse power law (IPL) PDF *ψ*(*τ*) ∝1/(*τ*)^*μ*^ with 1 < *μ* < 3. We establish the coupling between two temporally complex systems using a phenomenological approach inspired by models of swarm cognition and prove that complexity matching, namely sharing the same IPL index *μ*, facilitates the transport of information, generating perfect synchronization, reminiscent of, but distinct from chaos synchronization. This advanced form of complexity matching is expected to contribute a significant progress in understanding and improving the bio-feedback therapies.

**Author Summary:** This paper is devoted to the control of complex dynamical systems, inspired to real processes of biological and sociological interest. The concept of complexity we adopt focuses on the assumption that the processes of self-organization generate intermittent fluctuations and that the time distance between two consecutive fluctuations is described by a distribution density with an inverse power law structure making the second moment of these time distances diverge. These fluctuations are called crucial events and are responsible for the ergodicity breaking that is widely revealed by the experimental observation of biological dynamics. We argue that the information transport from one to another complex system is ruled by these crucial events and we propose an efficient theoretical prescription leading to qualitative agreement with experimental results, shedding light into the processes of social learning. The theory of this paper is expected to have important medical applications, such as an improvement of the biofeedback techniques, the heart-brain communication and a significant progress on cognition and the contribution of emotions to cognition.

## I. INTRODUCTION

It is slightly over a half century since Ross Ashby, in his masterful book [1], alerted scientists to be aware of the difficulty of regulating biological systems, and that “the main cause of difficulty is the variety in the disturbances that must be regulated against”. This insightful observation need not lead to the conclusion that complex systems cannot be regulated. It is possible to regulate them if the regulators share the same high intelligence (complexity) as the systems being regulated. Herein we refer to the Ashby’s *requisite variety* with the modern term *complexity matching* [2]. The term complexity matching has been widely used in the recent past [3–8] including the synchronization between the finger tapping and a complex metronome interpreted to be a system as complex as the human brain. These synchronization studies are today’s realizations of the regulation of the brain, in conformity with the observations of Ashby.

It is important to stress that there exists further research directed toward the foundation of social learning [9–12] that is even more closely connected to the ambitious challenge made by Ashby. In fact, this research aims at evaluating the transfer of information from the brain of one player to that of another, by way of the interaction the two players established through their avatars [10]. The results are exciting in that the trajectories of the two players turn out to be significantly synchronized. But even more important than synchronization is the fact that the trajectories of the two avatars have a universal structure based on the shared EEGs of the human brain.

This paper provides a theoretical understanding of the universal structure representing the brain of the two interacting individuals. In addition, the theory can be adapted to the communication, or information transfer, between the heart and the brain [13] of a single individual.

The transfer of information between interacting systems has been addressed using different theoretical tools, examples of which include: *chaos syncronization* [14], *self-organization* [15], and *resonance* [16]. On the other hand, in a system as complex as the brain [17] there is experimental evidence for the existence of crucial events. These crucial events, for our purposes here, can be interpreted as organization rearrangements, or renewal failures. The interval between consecutive crucial events is described by a waiting-time IPL PDF *ψ*(*t*) ∝1*/t*^*μ*^, with 1 < *μ* < 3. The crucial events generate ergodicity breaking and are widely studied to reveal fundamental biological statistical properties [18].

Another important property of biological processes is homeodynamics [19], which seems to be in conflict with homeostasis as understood and advocated by Ashby. Lloyd *et al* [19] invoke the existence of bifurcation points to explain the transition from homeostasis to homeodynamics. This transition, moving away from Ashby’s emphasis on the fundamental role of homeostasis, has been studied by Ikegami and Suzuki [20] and by Oka *et al*.[21] who coined the term *dynamic homeostasis*. They used Ashby’s cybernetics to deepen the concept of self and to establish if the behavior of the Internet is similar to that of the human brain.

Experimental results exist for the correlation between the dynamics of two distinct physiological systems [22], but they are not explained using any of the earlier mentioned theoretical approaches [14–18]. Herein we relate this correlation to the occurrence of crucial events. These crucial events are responsible for the generation of 1*/f* noise, *S*(*f*) ∝1*/f* ^3*-μ*^ [23] and the results of the psychological experiment of Correll [24]. The experimental data imply that activating cognition has the effect of making the IPL index *μ* < 3 cross the barrier between the Lévy and Gauss basins of attraction, namely making *μ >* 3 [25]. This is in line with Heidegger’s phenomenology [26].

The crossing of a basin’s boundary is a manifestation of the devastating effect of violating the linear response condition, according to which a perturbation should be sufficiently weak as to not affect a system’s dynamic complexity [27]. The experimental observation obliged us to go beyond the linear response theory adopted in earlier works in order to explain the transfer of information from one complex system to another. This transfer was accomplished through the matching of the IPL index *μ* of the crucial events PDF of the regulator with the IPL index *μ* of the crucial events PDF of the system being regulated [28, 29]. This is consistent with the general idea of complexity matching [2] with the main limitation, though, that the perturbation intensity has to be sufficiently small as to make it possible to observe the influence of the perturbing system on the perturbed through ensemble averages, namely a mean over many realizations [28], or through time averages, if we know the occurrence time of crucial events [29].

Earlier work [30], based on the direct use of the dynamics of two complex networks, studied the case when a small fraction of the units of the driven system peceive the mean field of the driving system. At criticality the choice made by these units is interpreted as *swarm intelligence* [31], and, in the case of the Decision Making Model (DMM) adopted in [30] is associated to the index *μ* = 1.5. In [30] this synchronization is observed when both systems are in the supercritical condition and it is destroyed if one system is in the subcritical regime and the other in the supercritical regime, or viceversa. This suggests that the maximal synchronization is realized when both systems are at criticality, namely, they share the same power law index *μ*.

This ideal condition of complexity matching is studied in this paper supplemented by homeodynamics, not considered in the earlier work. The theory that we use herein should not be confused with the unrelated phenomenon of chaos synchronization. In fact, the intent of the present approach is to establish the proper theoretical framework to explain, for instance, brain-heart communication. Here the heart is considered to be a complex, but not chaotic, system, in accordance with a growing consensus that the heart dynamics are not chaotic [32].

## II. METHOD

To deal with the ambitious challenge of Ashby [1] we adopt the perspective of subordination theory [33]. This theoretical perspective is closely connected to the Continuous Time Random Walk (CTRW) [34, 35], which is known to generate anomalous diffusion. We use this viewpoint to establish an approach to explaining the experimental results showing the remarkable oscillatory synchronization between different areas of the brain [22].

Consider a clock, whose discrete hand motion is punctuated by ticks and the time interval between consecutive ticks is, Δ*t* = 1, by assumption. At any tick the angle *θ* of the clock hand increases by 2*π/T*, where *T* is the number of ticks necessary to make a complete rotation of 2*π*. We implement subordination theory by selecting for the time interval between consecutive ticks a value *τ/ < τ >* where *τ* is picked from an IPL waiting-time PDF *ψ*(*τ*) with a complexity index *μ >* 2. This is a way of embedding crucial events within the periodic process. Notice that in the Poisson limit *μ → ∞* the resulting rotation becomes virtually indistinguishable from those of the non-subordinated clock. Note further, that when *μ >* 2, the mean waiting time ‹*τ*› is finite. As a consequence, if *T* is the information about the frequency Ω = 2*π/T*, this information is not completely lost in the subordinated time series.

During the dynamical process the signal frequency fluctuates around Ω and the average frequency is changed into an effective value

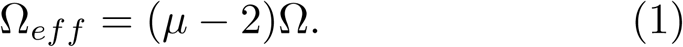

This formula can be easily explained with no need of going through a rigorous demonstration [36]. In fact, *μ* = 3 is the border with the Gaussian region *μ >* 3 where both the first and second moment of *ψ*(*τ*) are finite, and the average of the fluctuating frequencies is identical to Ω. In the region *μ* < 2 the process is non-ergodic, the first moment < *τ >* is divergent and the direct indications of homeodynamics vanish. The condition 2 < *μ* < 3 is compatible with the emergence of a stationary correlation function, in the long-time limit, with *μ* replaced by *μ* −1. Thus, using the result of earlier work [37] we obtain for the equilibrium correlation function exponentially damped regular oscillations. At the end of this oscillatory process, an IPL tail proportional to 1*/t*^*μ*-^1is obtained. Using a Tauberian theorm explains why the power spectrum *S*(*f*) becomes proportional to 1*/f* 3^*-μ*^ for *f →* 0. In summary, in a log-log representation, we obtain a curve with different slopes, *β* = 3- *μ*, to the left of the frequency-generated bump, and *β* = 2, to its right. The slope *β* = 2 is a consequence of the exponentially damped oscillations. Fig. (1) illustrates the result of a numerical approach to subordination, confirming the theoretical predictions.

**FIG. 1:**
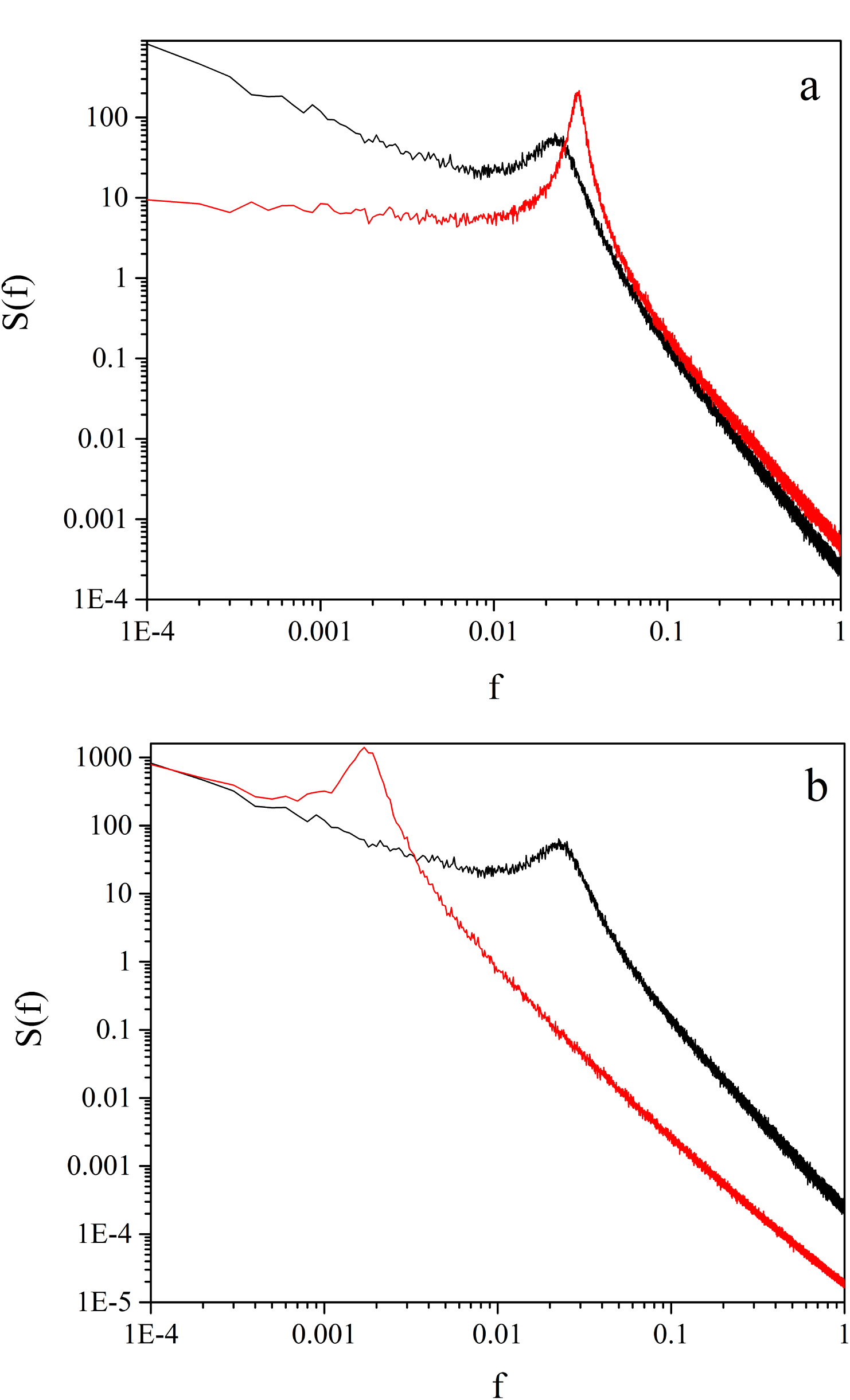
The spectrum *S*(*f*) of subordinations to the regular clock motion. (a) Ω = 0.06283, *μ* = 2.1 (black curve), *μ* = 2.9 (red curve). (b) *μ* = 2.1, Ω = 0.06283 (black curve), Ω = 0.0006283 (red curve).

We interpret the time evolution of *x*(*t*), the *x*- component of subordination to periodicity, as the result of a cooperative interaction between many oscillators. The IPL index *μ* quantifies the temporal complexity, spontaneously realized as an effect of oscillator-oscillator interactions. To make system-1 (S_1_) drive system-2 (S_2_) we have to generalize the swarm intelligence prescription adopted in earlier work [30, 31]. This generalization is necessary because the earlier work was based on the assumption that the single units of the complex systems, in the absence of interaction, undergo dichotomous fluctuations without the periodicity imposed here. In the absence of periodicity, the mean field *x*(*t*) of the complex system can be written as *x*(*t*) = (*U* (*t*) − *D*(*t*))*/N*, where *U* (*t*) is the number of individual in the state “*Up*”, *x >* 0, and *D*(*t*) is the number of individuals in the state “*Down*”, *x* < 0. In the case considered herein the number of units in a system *N* = *U* (*t*)+*D*(*t*) is constant. Using this notation (see Section III) we show that system-2 under influence of system-1 changes as:

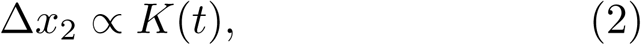

Where

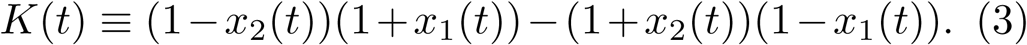

To properly take periodicity into account note that the mean field in S_1_ given by *x*_1_(*t*) has the functional form

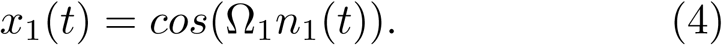

The mean field in S_2_ has the same periodic functional form, up to a time-dependent phase,

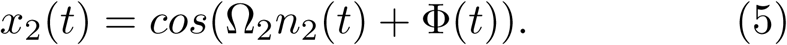

The phase Φ(*t*) is a consequence of the fact that the units of S_2_ try to compensate for the effects produced by the two independent self-organization processes. The number of ticks of the S_1_ clock, *n*_1_(*t*), due to the occurrence of crucial events, becomes increasingly different from the number of ticks of the S_2_ clock, *n*_2_(*t*). The units of S_2_, try to imitate the choices made by the units of S_1_. This is modeled by adjusting the phase Φ(*t*) of Eq. (5). The phase change is proportional to *K*(*t*) and to the derivative of *x*_2_(*t*) with respect to t. Thus, we obtain the central algorithmic prescription of this paper:

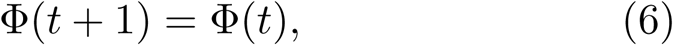

if at *t* + 1 no crucial event occurs, and

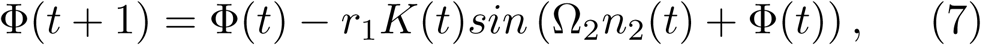

if at *t* + 1 a crucial event occurs. Note that the real positive number *r*_1_ *<* 1, defines the proportionality factor left open by Eq. (2), or, equivalently, defines the strength of the perturbation that S_1_ exerts on S_2_.

Fig. (2) illustrates the significant synchronization between the driven and the driving system obtained for *μ* = 2.2, close to the values of the crucial events of the brain dynamics [17]. This result can be used to explain the experimental observation of the synchronization of two people walking together [3] (see Section III).

The top panel of Fig. (3) shows that S_2_, with *μ*_2_ = 2.9, very close the Gaussian border, adopts the higher complexity of S_1_ with *μ*_1_ = 2.1, namely the complexity of a system very close to the ideal condition, *μ* = 2, to realize 1*/f* noise.

In the bottom panel of Fig. (3) we see that a driving system very close to the Gaussian border does not make the driven system less complex, but it does succeed in forcing it to adopt the regulator’s periodicity. Here we have to stress that the perturbing system is quite different from the external fluctuation that was originally adopted to mimic the effort generated by a difficult task [24, 25]. In that case, according to Heidegger’s phenomenology [26] the transition from *ready-to-hand* to *unready-to-hand* makes the IPL index *μ* depart from the 1*/f*-noise condition *μ* = 2 [24, 25] so as to reach the Gaussian border *μ* = 3 and to go beyond it.

The theory developed herein substantiates the opposite effect of cooperation. It is straightforward to extend this treatment to the case where S_1_ is influenced by S_2_ in the same way S_2_ is influenced by S_1_. To make this extension we have to introduce the new parameter *r*_2_, which defines the intensity of the influence of S_2_ on S_1_. As a result of this mutual interaction, we have 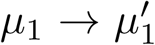 and 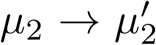. When *μ*_1_ < *μ*_2_ we expect

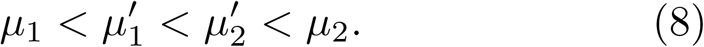

Fig. 4 shows that 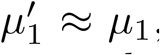 *≈μ*_1_, thereby suggesting that the system with higher complexity does not perceive its interaction with the other system as a difficult task, which would force it to increase its own *μ* [24, 25], while the less complex system has a sense of relief. We interpret this result as an important property that should be the subject of psychological experiments to shed light on the mechanisms facilitating the teaching and learning process. The theory underlying *complexity matching* and therefore requisite variety, makes it possible to go beyond the limitation of the earlier work on complexity management, as illustrated in Section III.

The term “intelligent” that we are using herein is equivalent to assessing a system to be as close as possible to the ideal condition *μ* = 2, corresponding to the ideal 1*/f* noise. In this sense two very intelligent systems are the brain and heart that when healthy share the property of a *μ* being close to 2. This paper therefore provides a rationale for (an explanation of) the synchronization between heart and brain time series [13] and shows that the concept of resonance, based on tuning the frequency of the stimulus to that of the system being perturbed, may not be appropriate for complex biological systems. Resonance is more appropriate for a physical system, where the tuning has been adopted over the years for the transport of energy not information. The widely used therapies resting on bio-feedback [38], are the subject of appraisal [39] and the present results may contribute to making therapeutic progress by establishing their proper use.

## III. MATERIAL, METHODS AND SUPPORTING INFORMATION

The main aim of this section is to afford an example of the experimental data that the theory of this paper can properly analyze. We focus on the close connection between Fig. 2 of this paper and Fig. 3 of Ref. [40]. For reader’s convenience we illustrate this connection with the help of Fig. 5.

**FIG. 2:**
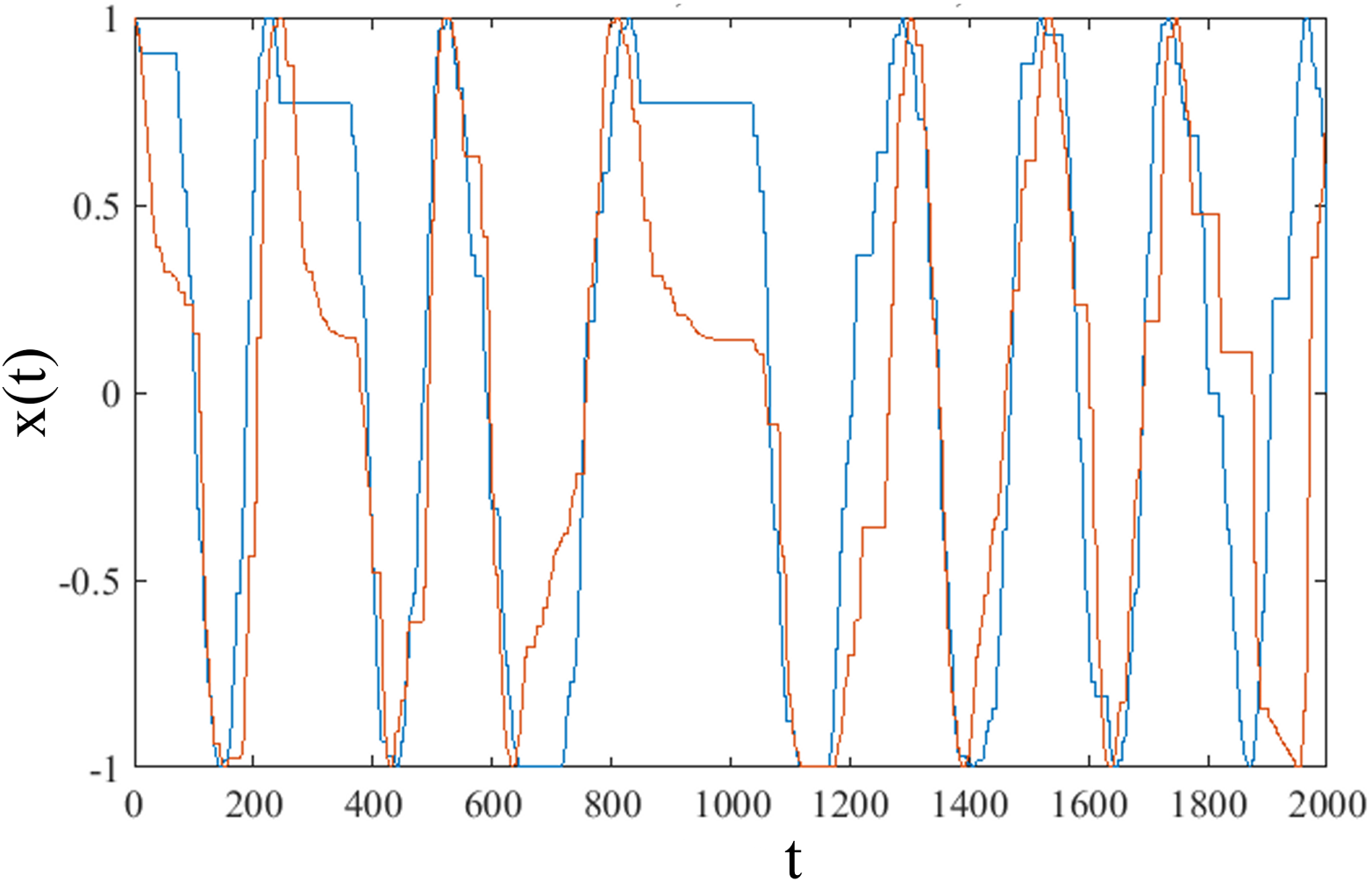
S_1_ (blue curve) drives S_2_ (red curve). Two systems are identical: *μ* = 2.2, Ω = 0.063, *r*_1_ = 0.05. The connection (one directional) is realized using Eq.(2) and Eq. (14).

**FIG. 3:**
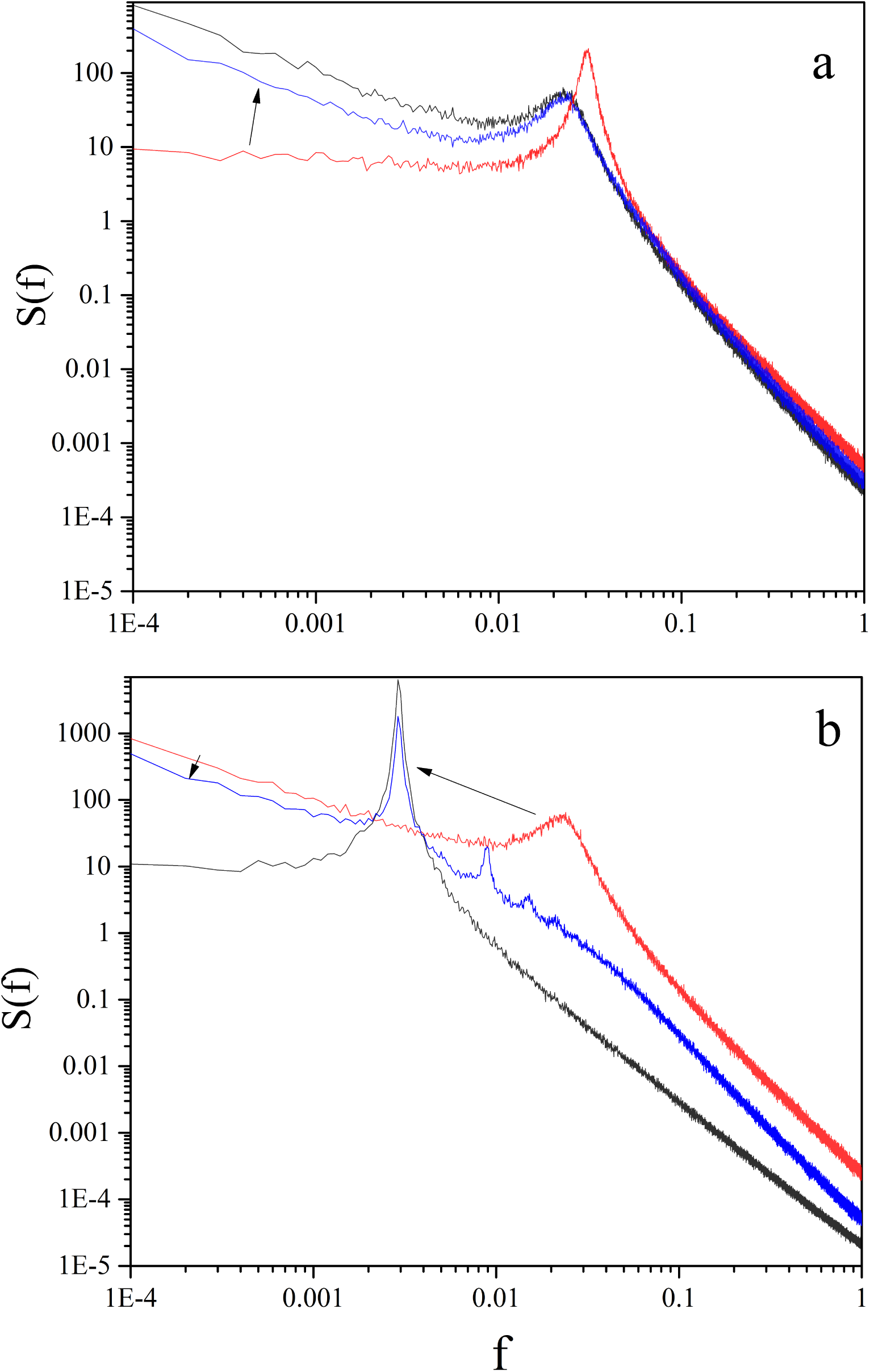
The spectrums of subordinations. (a) Black curve (*S*_1_): *μ* = 2.1, Ω = 0.063; red curve (*S*_2_):*μ* = 2.9, Ω = 0.063; blue curve: *S*_2_ after being connected (one directional) to *S*_1_ with *r*_1_ = 0.1. (b) Black curve (*S*_1_): *μ* = 2.9, Ω = 0.0063; red curve (*S*_2_): *μ* = 2.1,Ω = 0.063; blue curve: *S*_2_ after being connected (one directional) to *S*_1_ with *r*_1_ = 0.1.

**FIG. 4:**
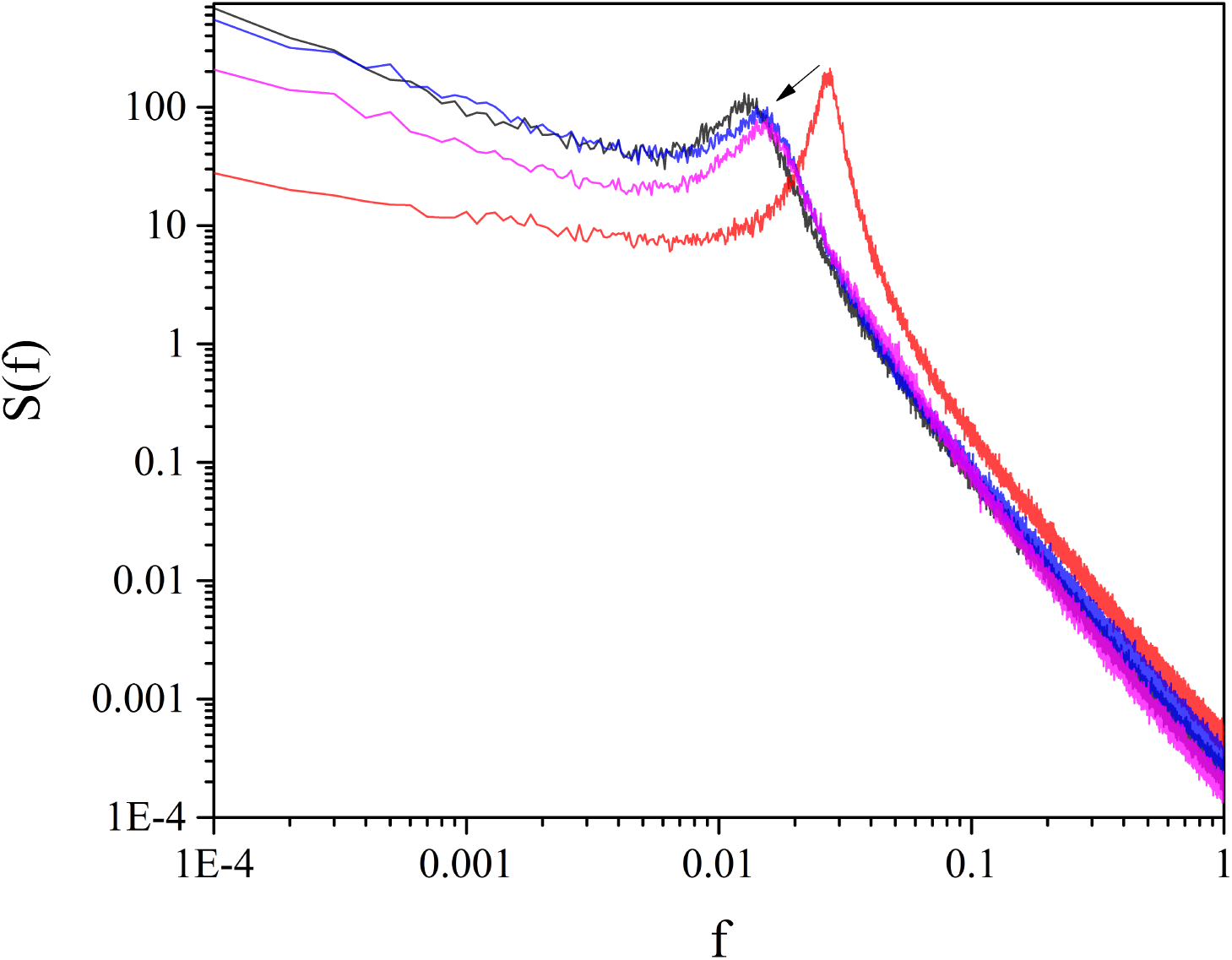
The spectrums of subordinations. Black curve (*S*1): *μ*_1_ = 2.2, Ω_1_ = 0.063; red curve (*S*_2_): *μ*_2_ = 2.9, Ω_2_ = 0.063. Blue and pink curves are the spectrums of *S*_1_ and *S*_2_ after being connected (bidirectional) with *r*_1_ = *r*_2_ = 0.1 respectively.

**FIG. 5:**
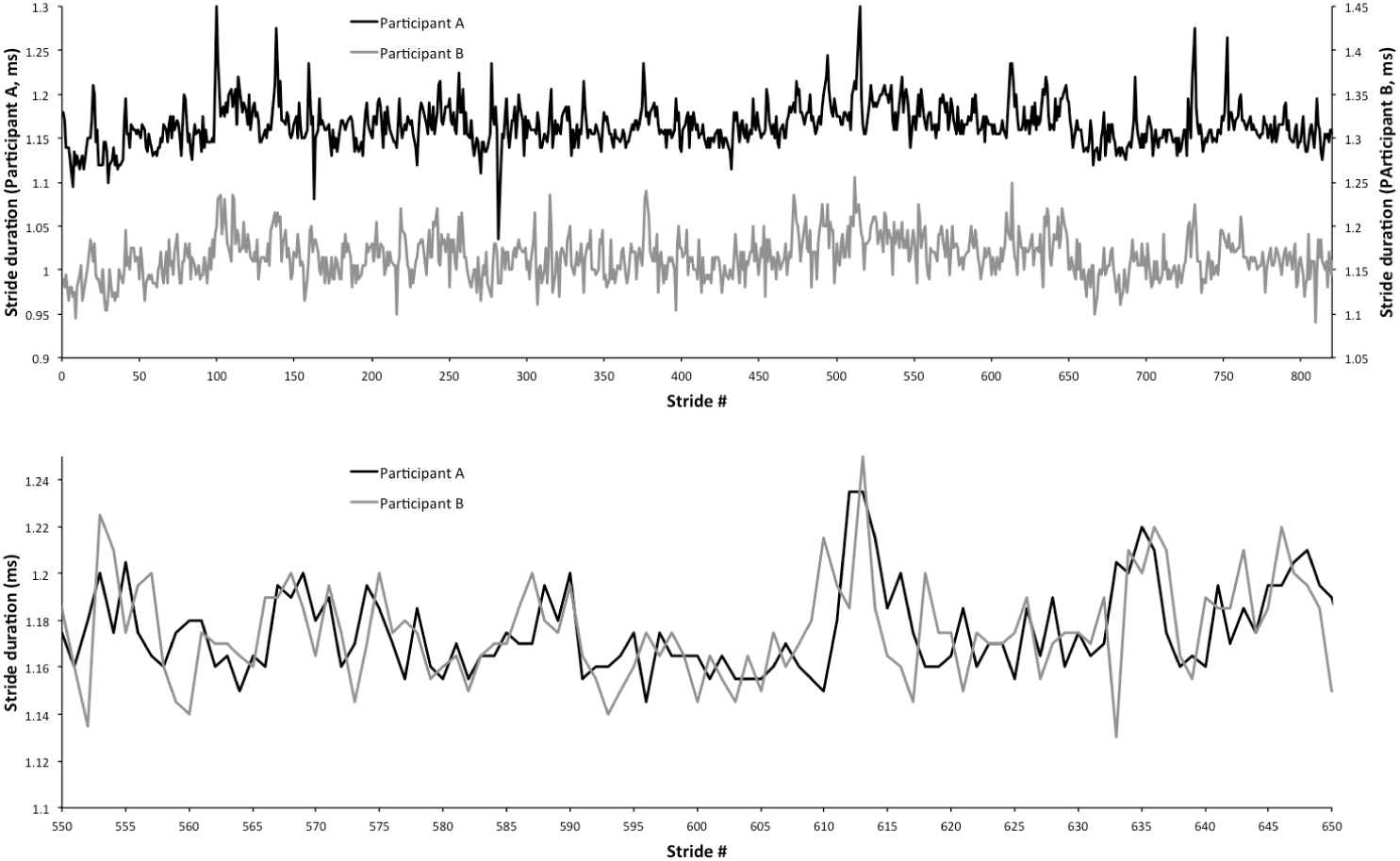
Experimental walking synchronization. These results have been derived with permission from Ref. [40]. The top panel shows two distinct walking trajectories. These are two human subjects trying to walk together. The bottom panel shows the same trajectories so as to emphasize their synchronization.

To explain how to derive the experimental synchronization we use Eqs. (2-5) of the text and we afford details on how to derive these important equations.

We use numerical results of the same kind as those illustrated in Fig. 2 of the text, properly modified to connect the two trajectories back to back. To make the qualitative similarity with the results of the experiment of Ref. [40] more evident we adopt the same prescription as that used by Deligniéres and his co-workers. We interpret the time distance between two consecutive crossings of the origin, *x* = 0, of Fig. 2 of the text as the time duration of a stride. We evaluate the mean stride duration and for the driven and the driving, for any stride we plot the deviation from this mean value. Fig. 6 illustrates the result of this procedure. This procedure can also be used to explain the synchronization between heart and brain [41].

**FIG. 6:**
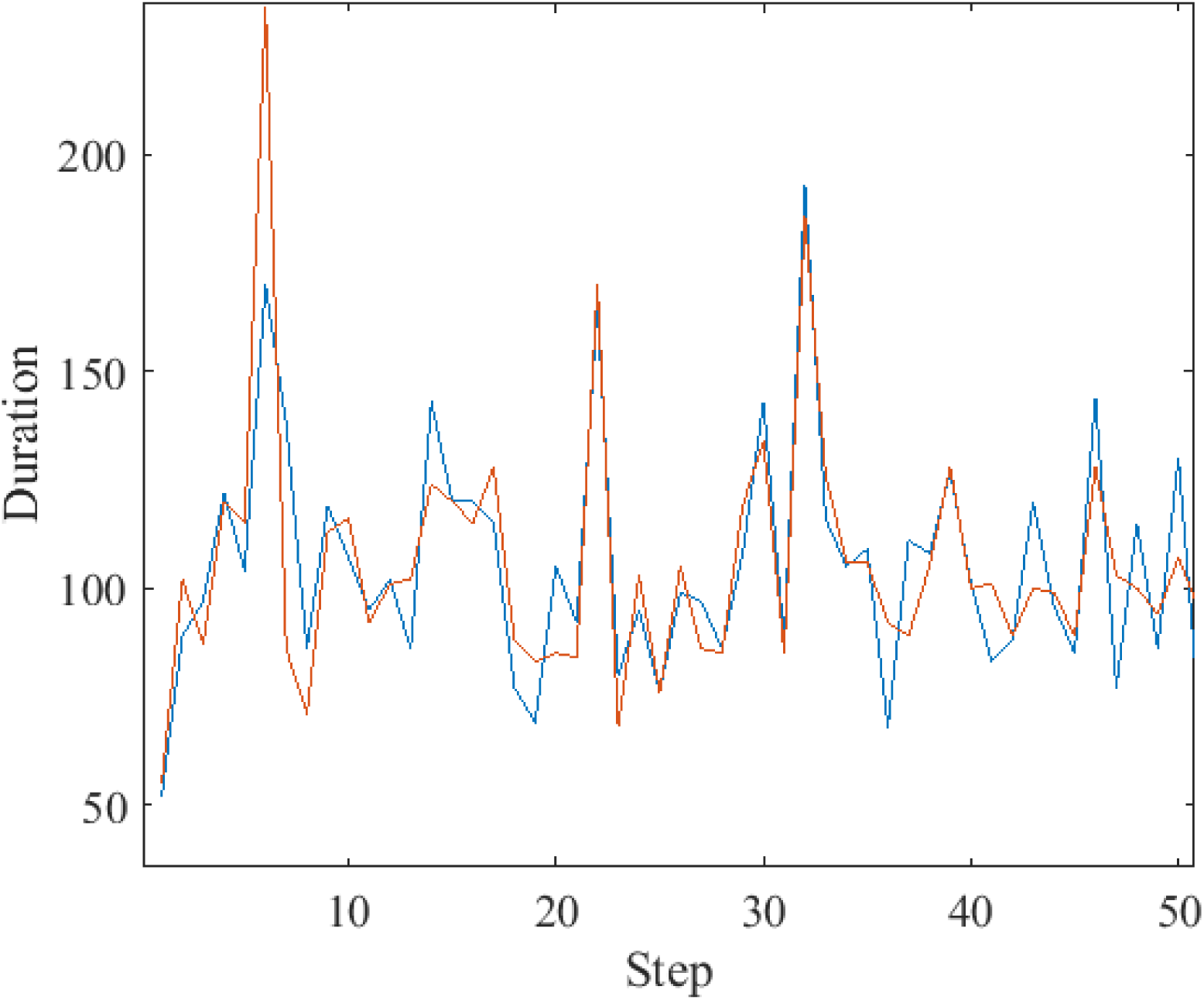
Time difference between the events of two identical systems connected back to back,*μ*_1_ = *μ*_2_ = 2.2, Ω_1_ = Ω_2_ = 0.062, *r*_1_ = *r*_2_ = 0.1.

In addition to showing with the help of Figs. 5 and 6 how the method of this paper works on the experimental data, we also illustrate the new theory in action to evaluate the cross-correlation between the driven and the driving complex network, going beyond the limitations of the research work on complexity management [28, 29].

### A. Group intelligence

Although subordination theory does not explicitly depend on the interaction between different units with their own periodicity, the system S_2_ with coordinate *x*_2_(*t*) is driven by the system S_1_ with coordinate *x*_1_(*t*) by a prescription inspired to create a swarm intelligence [31] using the (DMM) [42]. At a given time the units of the driven systems look at the driving system and according to its state increase or decrease the phase of the driven system. We note that

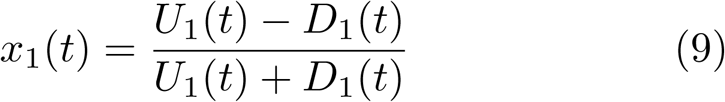

and

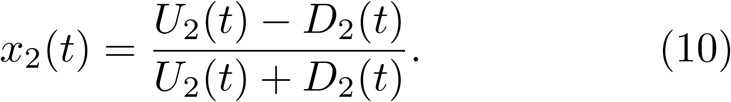

The symbols *U* and *D* denote the number of units in state “Up” and “Down”, respectively. The theory that we are using, detailed in Section II is based on subordination to periodicity. The single individuals of the complex system may have only the value 1, cooperation, or 1, defection. We introduce the angle *θ* to take periodicity into account and we interpret *cosθ* as the ratio of the difference between the number of cooperators and the number of defectors to the total number of units. Thus the majority of cooperators corresponds to 0 < *θ* < *π* and the majority of defectors corresponds to *π* < *θ* < 2*π*. We assume that the change in S_2_ because of interaction with S_1_ is

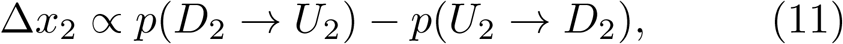

where the probability of making a transition from the state down to the state up in S_2_ is given by

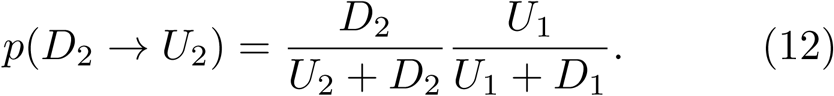

The form of Eq. (12) is due to the fact that this probability is the product of the probability of finding a unit in the driven system in the down state by the probability of finding a unit in the driving system in the up state. Using the same arguments we find

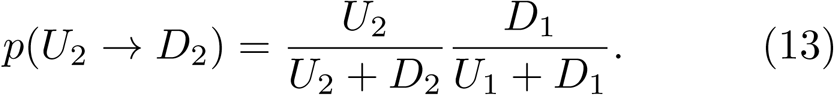

Let us plug Eq. (12) and Eq. (13) into Eq.(11). We obtain

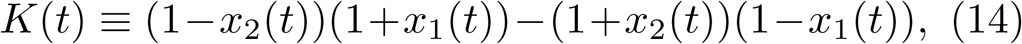

which is the important prescription of Eq. (3).

### B. Walking together

To facilitate appreciation of the similarity between the complexity matching prescription of this paper and the walking synchronization of Ref. [40], we invite the readers to look at the experimental results of Fig. 5. The real data are not available to us, and we use surrogate data instead. These surrogate data are derived from the numerical results adopted to get Fig. 6, with two ma-jor adjustmentsm, as earlier explained, of focusing on the observation of the origin crossing. The remarkably good qualitative agreement between Fig. (6) and Fig.(5) proves the efficiency of the complexity matching approach of this paper.

### C. Beyond Complexity Management

Complexity management is very difficult to observe. It is based on ensemble averages, thereby requiring the average over many identical realizations [28]. In the case of experimental signals of physiological interest, for instance on the brain dynamics, the ensemble average is not possible and recently a procedure was proposed [29] to convert an individual time series into many independent sequences, so as to have recourse again to an average over many realizations. This procedure, however, requires the knowledge of time occurrence of crucial events. The theory of this paper makes it possible to evaluate the correlation between the driving and the driven system using only one realization. We have to stress that while complexity management [28] does not affect the power index *μ* of the interacting complex networks, the theory of this paper, as shown by Fig. 3, affords the important information of how the cooperative interaction makes the unperturbed values of *μ* change as an effect of interaction.

For reader’s convenience, we illustrate the maximum value of the cross correlation function, *C*_*max*_, between the driving and the driven system in only two paradigmatic cases. Fig. 7 shows that a significantly large frequency difference strongly reduces the intensity of synchronization.

**FIG. 7:**
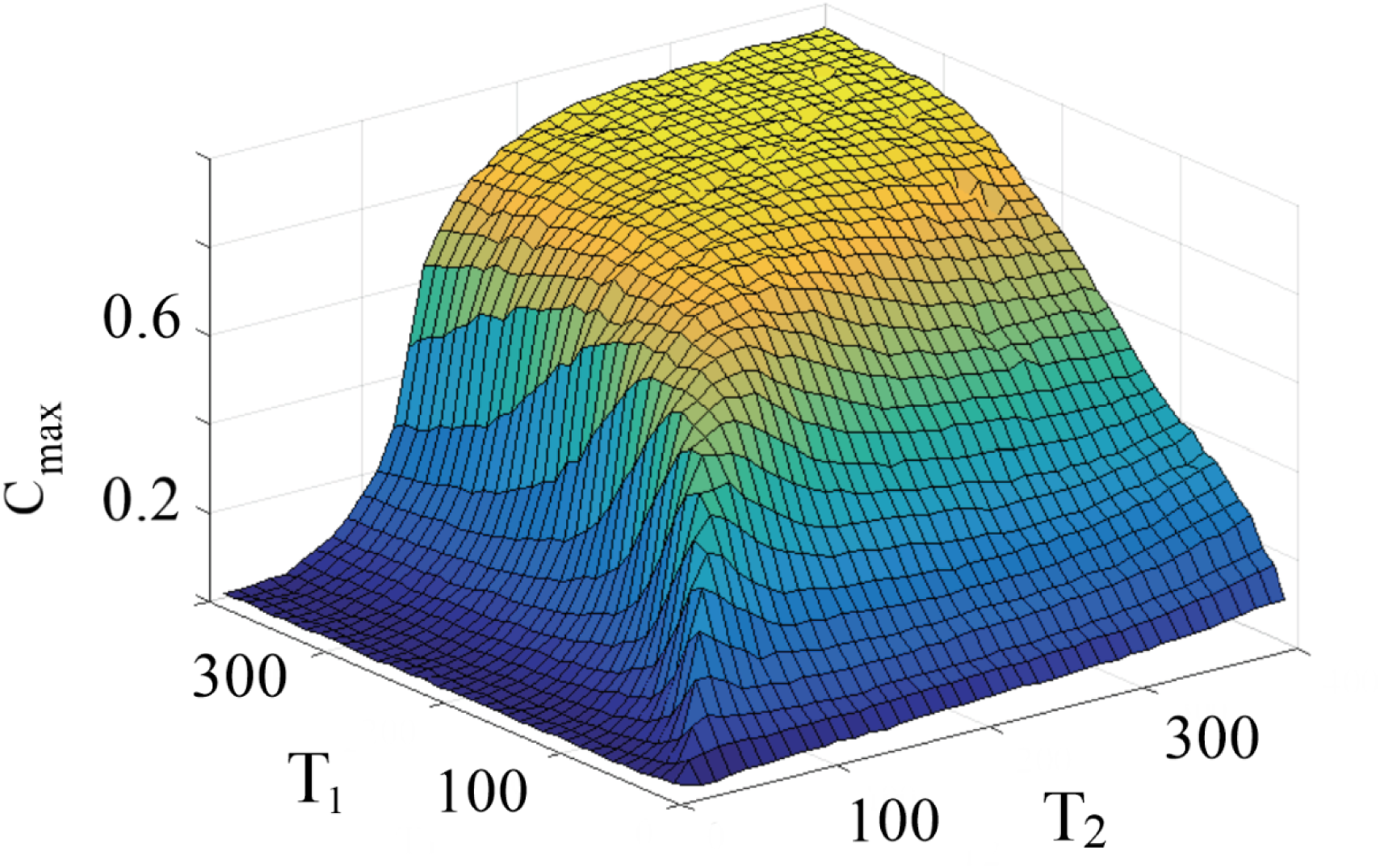
Dependence of *C*_*max*_ (as a measure for complexity matching) on the periodicity of the drive and driven systems. *μ*_1_ = *μ*_2_ = 2.8. *r*_1_ = 0.1.

Fig. 8 shows the effect of changing *μ*_1_ and *μ*_2_ on *C*_*max*_, while keeping the frequencies Ω identical.

**FIG. 8:**
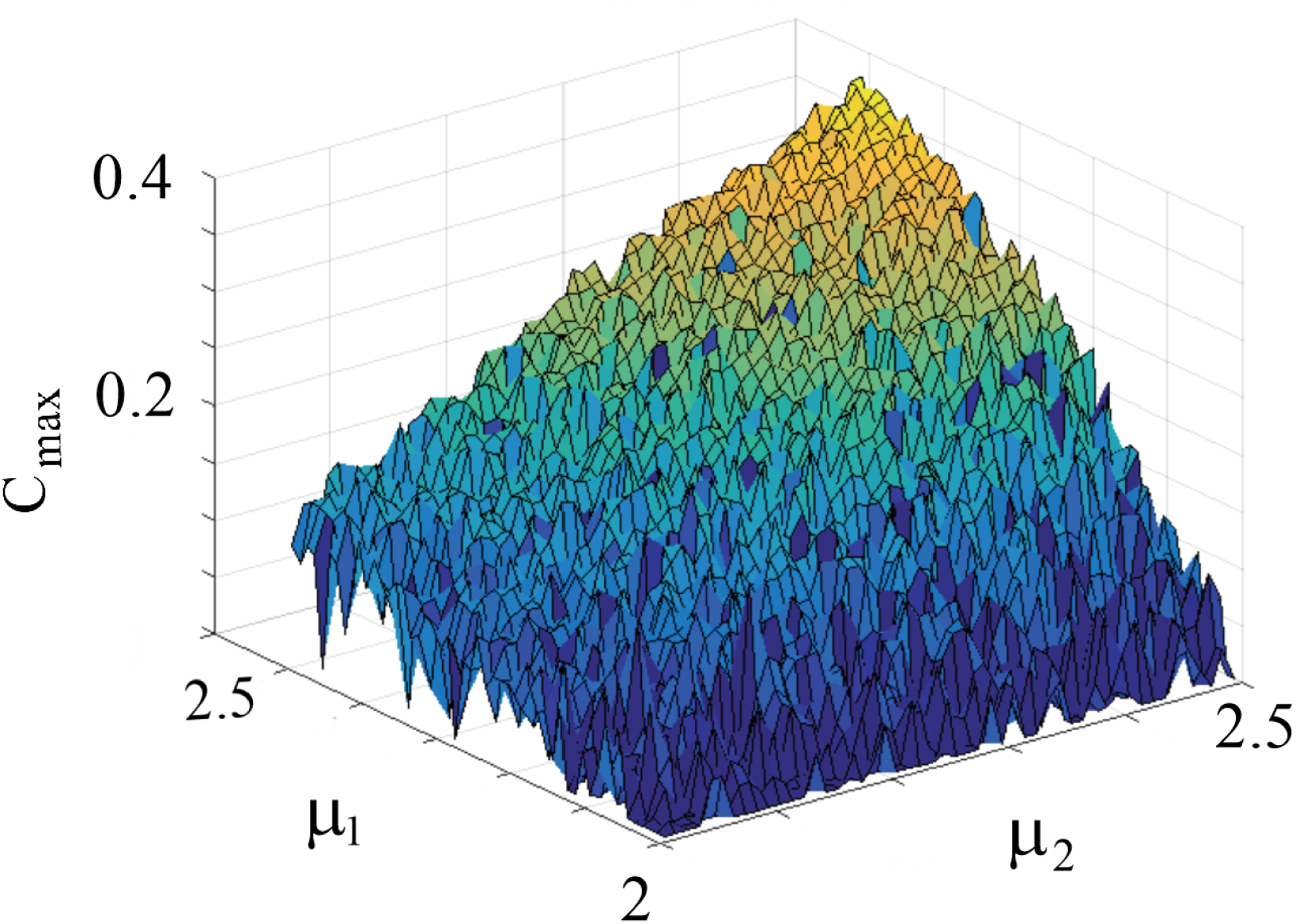
Dependence of *C*_*max*_ on the complexity index of the drive and driven systems. *T*_1_ = *T*_2_ = 50, *r*_1_ = 0.1.

